# Study on the mechanism of HE4 in hyperoxia-induced alveolar damage in rats

**DOI:** 10.1101/2022.09.22.509120

**Authors:** Xiaofang Yan, Xing Feng, Yan Gao, Dawei Liu, Lin Bai, Lu Xu

## Abstract

**Objective:** This study investigated the mechanism of human epididymis protein 4 (HE4) in hyperoxia-induced alveolar damage in rats.

**Methods:** The pathological changes of SD rat lung tissue were detected by hematoxylin-eosin staining. Plasma protein levels were measured by enzyme-linked immunosorbent assay (ELISA). The expression of mRNA was detected by real-time RT-PCR.

**Results:** Hyperoxia intervention within 7 days could induce acute lung injury in neonatal SD rats. Hyperoxia induction can increase HE4, MMP9 and TIMP1 in plasma and tissue of neonatal SD rats. By overexpressing and silencing HE4 gene in neonatal rat primary alveolar type II epithelial cells, we found that HE4 protein activates the phosphorylation of ERK and p65, activates the downstream MMP9 signaling pathway, inhibits MMP9 mRNA expression, inhibits protein activity, and reduces COLI degradation, increases collagen secretion, promoting matrix remodeling and fibrosis.

**Conclusion:** HE4 mediates the pathophysiological process of hyperoxia-induced lung injury in rats through ERK, MMP9 and TIMP1 signaling pathways. HE4 overexpression mediates lung basal remodeling and lung injury.

## Introduction

Bronchopulmonary dysplasia (BPD) is a common disease in the neonatal intensive care unit, which mostly occurs in preterm neonates of small gestational age and children with neonatal respiratory distress syndrome; it is caused by multiple factors in the perinatal period Due to premature and immature lung tissue, the result of acute lung injury and abnormal repair after injury can be complicated by different degrees of health problems, which brings great challenges to the family and society. With the improvement of neonatal treatment technology, the mortality rate of the disease has decreased, but the morbidity rate has not decreased significantly, and the smaller the gestational age and the lower the body weight, the higher the incidence of the disease; Therefore, how to identify early and find early predictive markers is a research hotspot.

In recent years, it has been studied that human epididymis protein 4 (HE4) is abnormally elevated in the plasma of children with neonatal respiratory dysfunction [1], and it is a new marker for predicting the prognosis of some pulmonary fibrosis diseases [2]. HE4 is a secreted glycoprotein that belongs to the WFDC domain family, a member of the antiprotease family consisting of the whey acidic protein (WAP) domain [3], and high expression of HE4 mRNA can be detected in the trachea and lung [4]. Knockout of this gene in animal studies can cause alveolar vascular dysplasia and severe dyspnea [5, 6]; and in pulmonary fibrosis diseases, HE4 interferes with pulmonary fibrosis by regulating matrix metalloproteinases (MMPs) signaling pathway and the remodeling process of the alveolar extracellular matrix [3, 7–9]. The key link in the pathophysiological process of BPD is acute lung injury and abnormal remodeling of the matrix after injury. Like pulmonary fibrosis, there are lung damage and remodeling. Therefore, it is reasonable to speculate that HE4 plays an important role in BPD.

The hyperoxia-induced BPD model is the most commonly used modeling method in current experimental research; the animals involved include rodents such as rats and mice. High-concentration oxygen exposure can lead to early acute lung injury in neonatal rats, and pulmonary fibrosis can gradually appear with the prolongation of high-concentration oxygen exposure. 85% oxygen concentration exposure for 7 days can lead to typical BPD-like manifestations such as blocked alveolar development, alveolar simplification, and disorder of angiogenesis [10]. Therefore, in this experiment, the hyperoxia-induced BPD rat animal model was selected, and the HE4 gene was overexpressed and silenced in the primary alveolar type II epithelial cells of neonatal rats to explore the mechanism of HE4 in the early stage of BPD, that is, acute alveolar injury. We speculate that HE4 mediates the process of lung basal remodeling and lung injury, and can be used as an early predictive marker for BPD.

## Experimental Materials and Methods

### Experimental materials

#### 1. Experimental animals

Neonatal SD rats were used in this experiment. This experiment was approved by the Experimental Animal Ethics Committee of our hospital (approval number 2020028), and the relevant experiments were carried out in accordance with the “3R” principle.

#### 2. Cell line

The neonatal mouse primary alveolar type II epithelial cell line was purchased from Wuhan Proceed Life Technology Co., Ltd. (Wuhan, China) and used throughout the study.

#### 3. Main reagents and antibodies

HE4, MMP9 and TIMP1 ELISA detection kits were purchased from Shanghai Enzyme Link Biotechnology; Anti-HE4 Antibody (Affinity, DF7703, 1:1000), Anti-KL-6 (Affinity, AF8524, 1:1000)) Anti-MMP9 Antibody (Affinity, GB12132-1, 1:1000), Anti-TIMP1 Antibody (Affinity, AF7007, 1:1000), Anti-Collagenl Antibody (Affinity, AF7001, 1:1000), Anti-ERK1/2 Antibody (Affinity, AF0155, 1:1000), Anti-Phospho-ERK1/2 Antibody (Affinity, AF1015, 1:1000), Anti-NF-κB Antibody (Affinity, AF5006, 1:1000), Anti-Phospho-NF-κB Antibody (Affinity, AF3389, 1:1000).

### Experimental method

#### 1. Experimental modeling and grouping

Neonatal SD rats (40) were randomly divided into normal control group and hyperoxia-induced group after birth. Each group was set at 3 time points of 24 hours, 3 days and 7 days. They were placed in air (20.8% oxygen concentration) and hyperoxia (85% oxygen concentration) environments for 7 days. During this period, nursing dams were housed with control and hyperoxia-exposed pups, and the dams were switched between room air and hyperoxia environments every 24 hours to avoid toxicity to the dams and to allow continued lactation. All animals were housed in a controlled environment with a 12-hour light/dark cycle and standard diet and water ad libitum. 3 days

#### 2. Rat lung tissue and blood collection

Five neonatal SD rats in each group were anesthetized by intraperitoneal injection of 5% chloral hydrate, blood was collected from the heart after anesthesia, and whole lungs were quickly collected on ice. After the heart was drawn, the blood was naturally coagulated for 20 min, centrifuged at 3000 rpm for 20 min, and the supernatant was collected and stored in a −80°C freezer. The left lung was fixed with 4% paraformaldehyde, embedded in paraffin after 48 hours, and stored in a refrigerator at 4°C. The right lung was frozen in liquid nitrogen and then transferred to a −80°C refrigerator for cryopreservation.

#### 3. Cell recovery, passage and grouping, and transfection

Neonatal murine primary alveolar type II epithelial cell lines were removed from liquid nitrogen. After resuscitation, the cells were cultured in a 5% CO2-based cell incubator with 10% fetal bovine serum at 37°C until the cells were in a stable state for subsequent experiments.

Construction and determination of recombinant plasmids: After the puro vector was digested with EcoR I and Hind 11I, it was digested with the same restriction enzymes as HE4 shDNA (HindIII-rat-HE4-F, 5’-GTTTAAACTTAAGCTTATGCCTGCTTGTCGCCTC-3’; EcoRI-rat-HE4-R 5 ‘-GATATCTGCAGAATTCTCAGAATTTGGGTGTGGTGCAG-3’) was ligated after annealing to construct recombinant plasmids targeting different sites. Continue to use E. coli containing pSUPER-HE4 shDNA. Coli DH5a was expanded and the plasmid was extracted and frozen at −80°C. The plasmid was successfully constructed using Lipo2000 to construct HE4 overexpressing cells, which were replaced with complete medium for 4 h. The experimental group was then injected with 3 L/min of high-purity gas mixture containing 85% oxygen, and then placed in a 50 mL/L CO2 incubator for 15 minutes, and incubated in 85% hyperoxia for 6 hours, while the control group was not treated with 85% oxygen. Cells were harvested after 6 hours of hyperoxia culture. HE4 gene was knocked out in cells using HE4 siRNA, and the cell culture method was consistent with HE4 overexpressing cells.

#### 4. ELISA detection

Take out the kit from the refrigerated environment and place it at room temperature for 15-30 minutes; take out the blood samples in advance to thaw, and operate in strict accordance with the kit instructions to calculate the sample concentration.

#### 5. HE staining

For hematoxylin and eosin (HE) staining, lung tissue was fixed, paraffin embedded, sectioned and stained with HE.

#### 6. Western blot

Take out the frozen lung tissue from the −80°C freezer, extract the protein and calculate the protein concentration after lysing, discard the culture medium from the cell sample, add 4°C pre-cooled PBS to wash the cells, and then aspirate and discard the culture. Put the culture plate after discarding PBS on ice, add 20 μl phosphatase inhibitor and 10 μl PMSF (100 mM) solution to 1 mL of lysis solution, shake well and place on ice for lysis, and calculate the protein concentration. Perform SDS-PAGE gel electrophoresis, calculate the current for membrane transfer by 1.5 mA/cm2 gel, and transfer the membrane to the solid-phase NC membrane in a semi-dry manner after 45 min of membrane transfer; After blocking at room temperature for 2 h, add the corresponding primary antibody, incubate overnight at 4°C, wash three times with TBST solution, add the corresponding secondary antibody, incubate at room temperature for 1 h, wash three times with TBST solution for 5 min each time, and expose to the chemiluminometer.

#### 7. RT-PCR

Total RNA extraction was carried out according to the steps of Trizol reagent. Trizol solution and chloroform were mixed 5:1 and left to stand for 5 minutes. Dissolve in free dH2O and measure the concentration; prepare reverse transcription reaction mixture on ice. The PCR primer sequences and amplified fragments are shown in Table 1 below. Pre-denaturation at 95°C for 5 min, followed by 30 cycles of 30 denaturation at 95°C, annealing at 60°C for 30 s, extension at 72°C for 30 s, and final extension at 72°C for 7 min. The mRNA expression of the gene of interest was normalized to cellular actin levels and analyzed for 2-ΔΔCt.

**Table 1:**
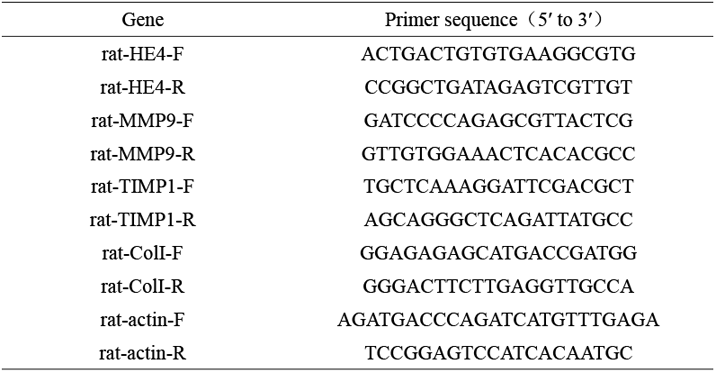
Base sequences of each primer

### Statistical analysis

Statistical analysis and graphing of all data in this study were achieved by SPSS 23.0 and Graphpad primsm7. For the description of normally distributed data, the mean ± standard deviation (±S) was used to describe, and the comparison between the two groups was performed by independent sample t test; the significance test level was 0.05.

## Results

### 1. HE staining results of two groups of lung tissues in neonatal SD rats induced by hyperoxia

In the normal control group, the alveolar structure was complete, and the thickness of the alveolar wall was uniform. Compared with the normal control group, the alveolar septa in the hyperoxia-induced group gradually became thinner and discontinuous, the alveolar wall was thin, the alveolar cavity gradually increased, some alveoli were fused, the number of alveoli was reduced, the size of the alveoli was different, and the alveolar structure was simplified. (figure 1)

**Figure 1:**
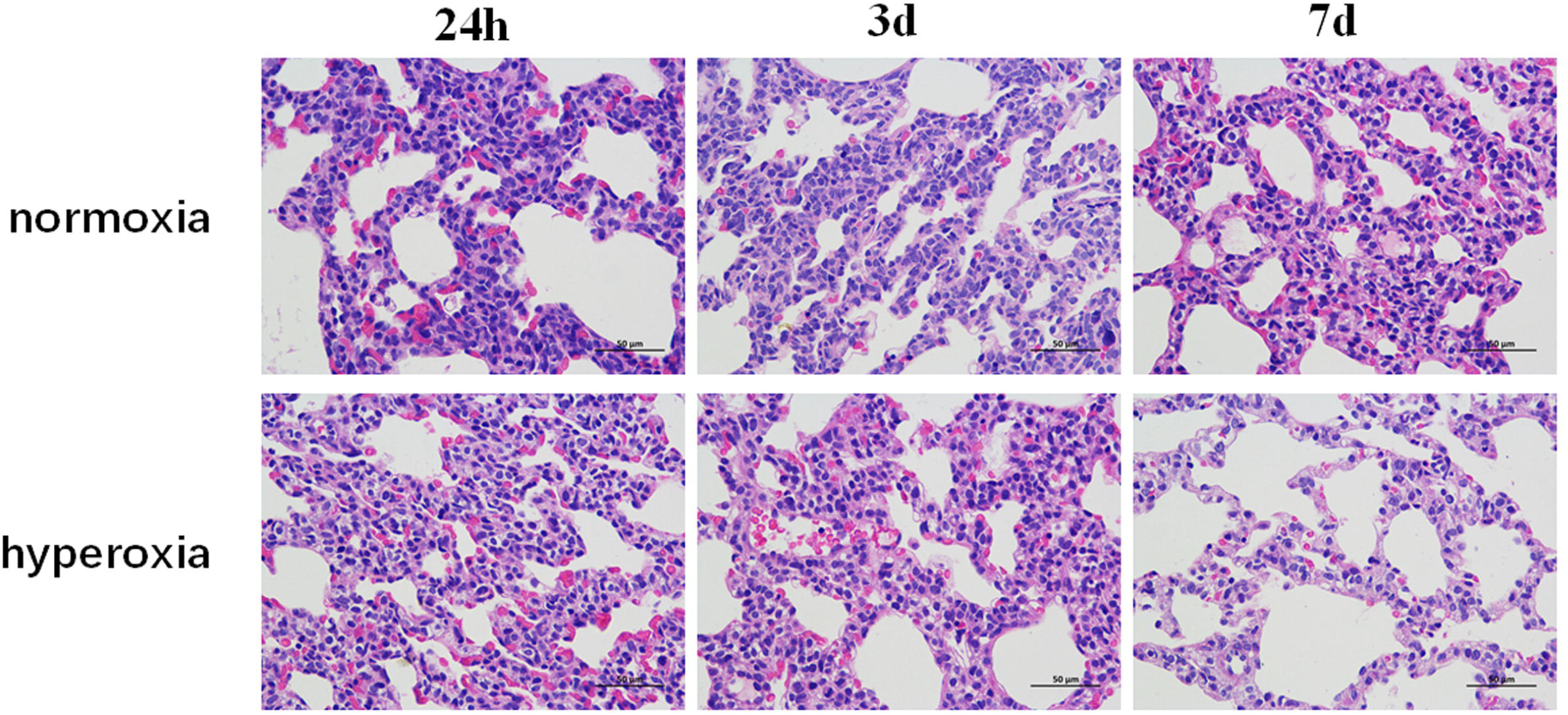
H&E(×400)

### 2. The results of ELSIA detection of HE4, MMP9 and TIMP1 protein concentrations in rat plasma

The protein levels of HE4, MMP9, and TIMP1 in the hyperoxia-induced group were significantly higher than those in the normal control group at 24 hours and 3 days, and the differences were statistically significant (p<0.05). Compared with the normal control group, the concentration of hyperoxia induction for 7 days was similar, and the difference was not statistically significant, p>0.05. (figure 2).

**Figure 2:**
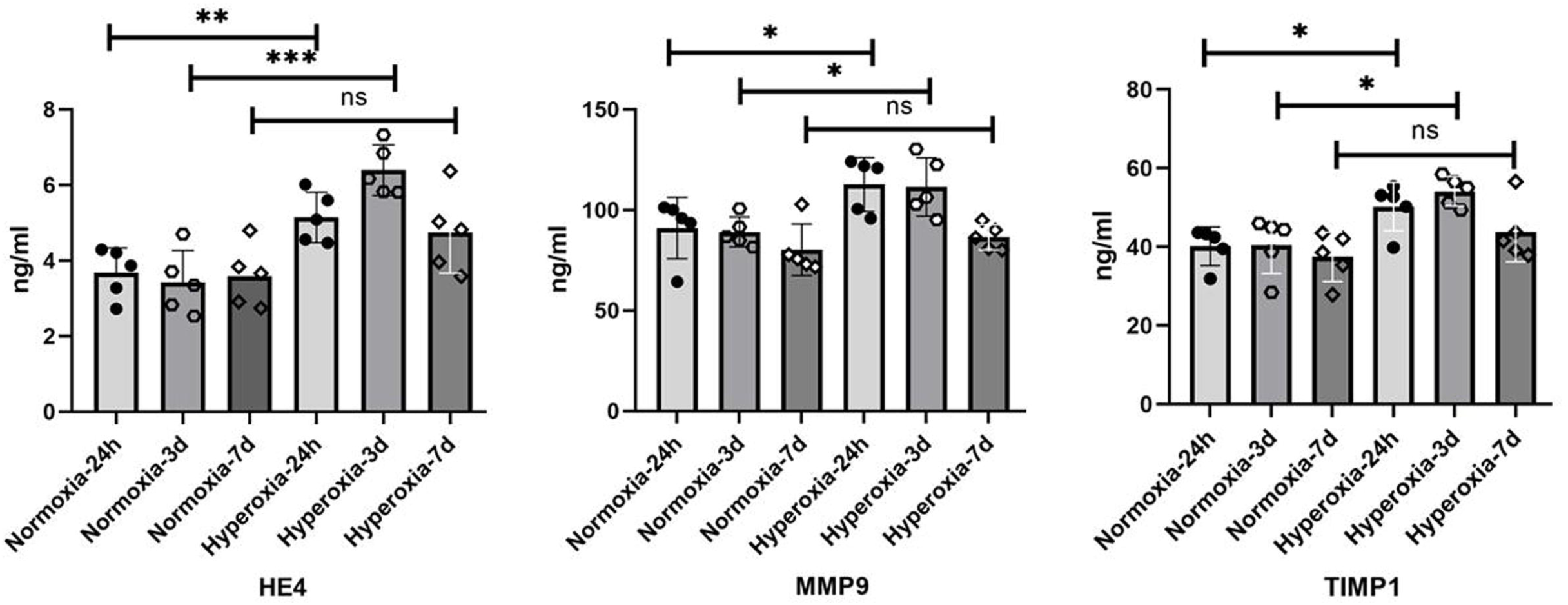
Changes of HE4, MMP9, TIMP1 concentrations in blood detected by ELSIA (mean±SEM, n=5) *p<0.05; **p<0.01; ***p<0.001; ns: no significant difference.

### 3. Western blot detection of HE4, MMP9 and TIMP1 protein expression in lung tissue

The expression level of HE4 protein in lung tissue was similar in normal control group at 24h, 3d, and 7d; The expression level of HE4 protein in the hyperoxia-induced group at 24 hours, 3 days and 7 days was higher than that of the normal control group at the corresponding time points. The expression levels of MMP9 protein in the normal control group were similar at 24h, 3 days and 7 days; The expression of MMP9 protein in the hyperoxia-induced group was higher than that in the normal control group at the corresponding time point, and the difference was statistically significant. The expression level of TIMP1 protein at each time point in the normal control group was similar; The expression of TIMP1 protein in the hyperoxia-induced group was higher than that in the normal control group at the corresponding time points, and the differences were statistically significant. (Figure 3)

**Figure 3:**
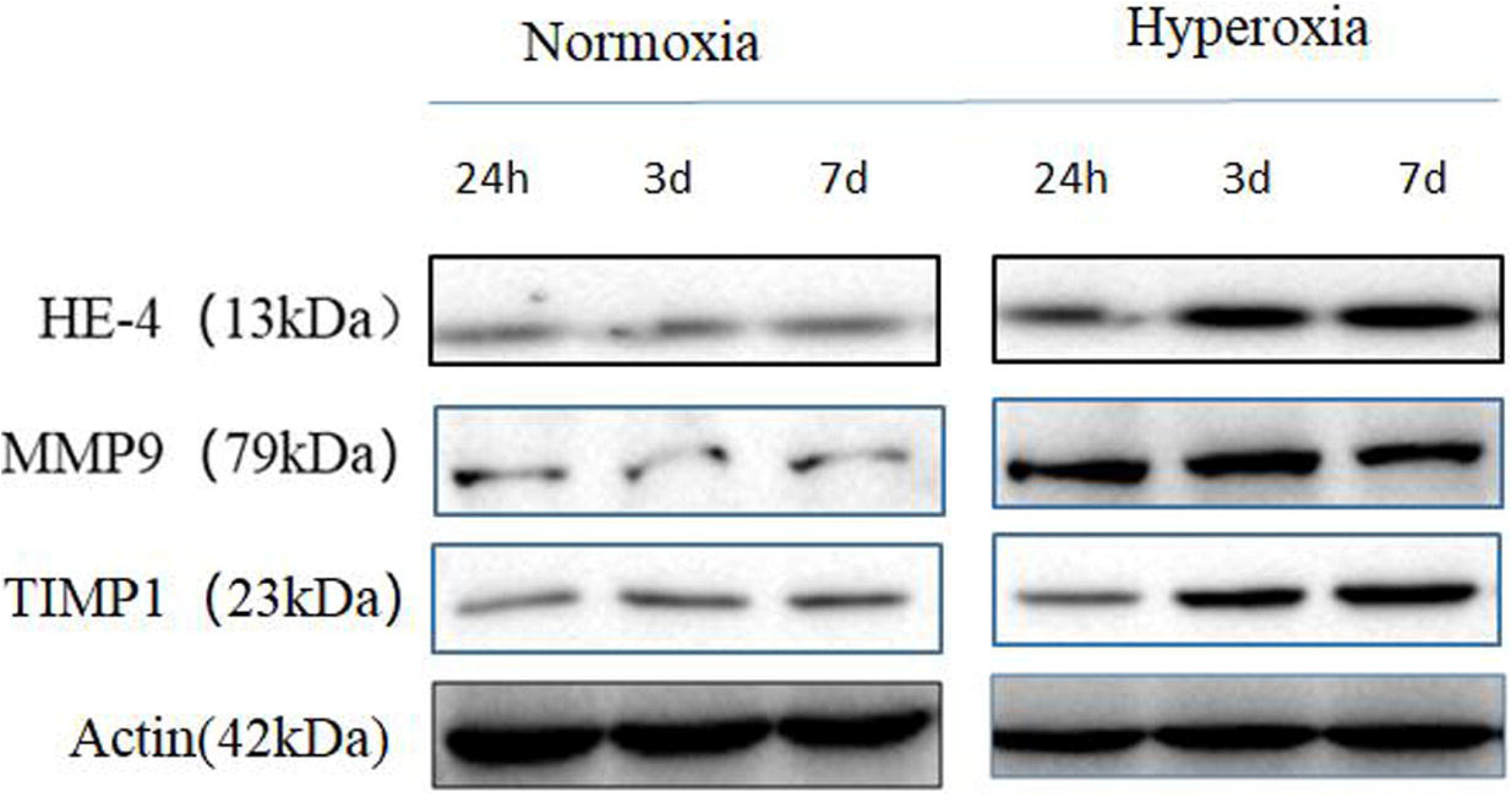
Western blot detection of HE4, MMP9, TIMP1 protein expression in lung tissue (n=5)

### 4. RT-PCR detection of HE4, MMP9 and TIMP1 mRNA expression in lung tissue

The expression of HE4, MMP9 and TIMP1 mRNA in lung tissue was significantly higher than that in the normal control group at 24 hours and 3 days after hyperoxia induction, and the difference was statistically significant (p<0.05). The expression of HE4 and TIMP1 mRNA in lung tissue was significantly higher than that of the normal control group at 7 days of hyperoxia induction, and the difference was statistically significant (p<0.05). The expression trends of HE4 and TIMP1 mRNA were consistent, and the expression was most significantly up-regulated when hyperoxia was induced for 3 days; The expression of MMP9 mRNA was most significantly up-regulated at 24 hours of hyperoxia intervention, and then the expression level gradually decreased. (Figure 4)

**Figure 4:**
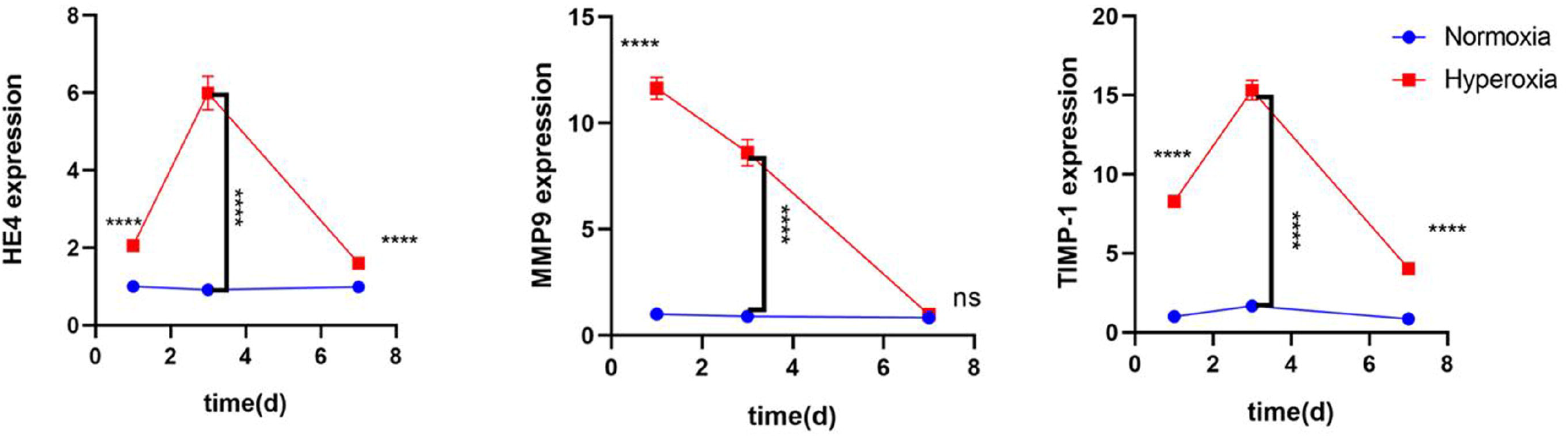
RT-PCR detection of the relative mRNA expression of HE4, MMP9, and TIMP1 in lung tissue (mean±SEM, n=5): ****p<0.0001; ns, no statistical difference

### 5. Western blot detection of target protein content in cell samples

The protein contents of HE4, TIMP1, COL1, p-ERK and p-p65 in the hyperoxia culture group were higher than those in the normal control group, but the MMP9 protein content was significantly lower than that in the normal control group, and the above differences were statistically significant (p<0.05); Compared with the normal control group, the MMP9 protein content in the gene overexpression group was significantly lower, while the TIMP1, COL1, p-ERK and p-p65 protein content were significantly higher than those in the normal control group, and the difference was statistically significant (t=46.0, p<0.05)); Compared with the normal control group, the protein level of MMP9 in the HE4 gene silencing group was significantly increased, and the protein levels of TIMP1 and COL1 were significantly decreased compared with the normal control group, and the above differences were statistically significant (p<0.05); However, the protein levels of p-ERK and p-p65 in the gene silencing group were similar to those in the normal control group, and there was no statistical significance (p>0.05). (Figure 5)

**Figure 5:**
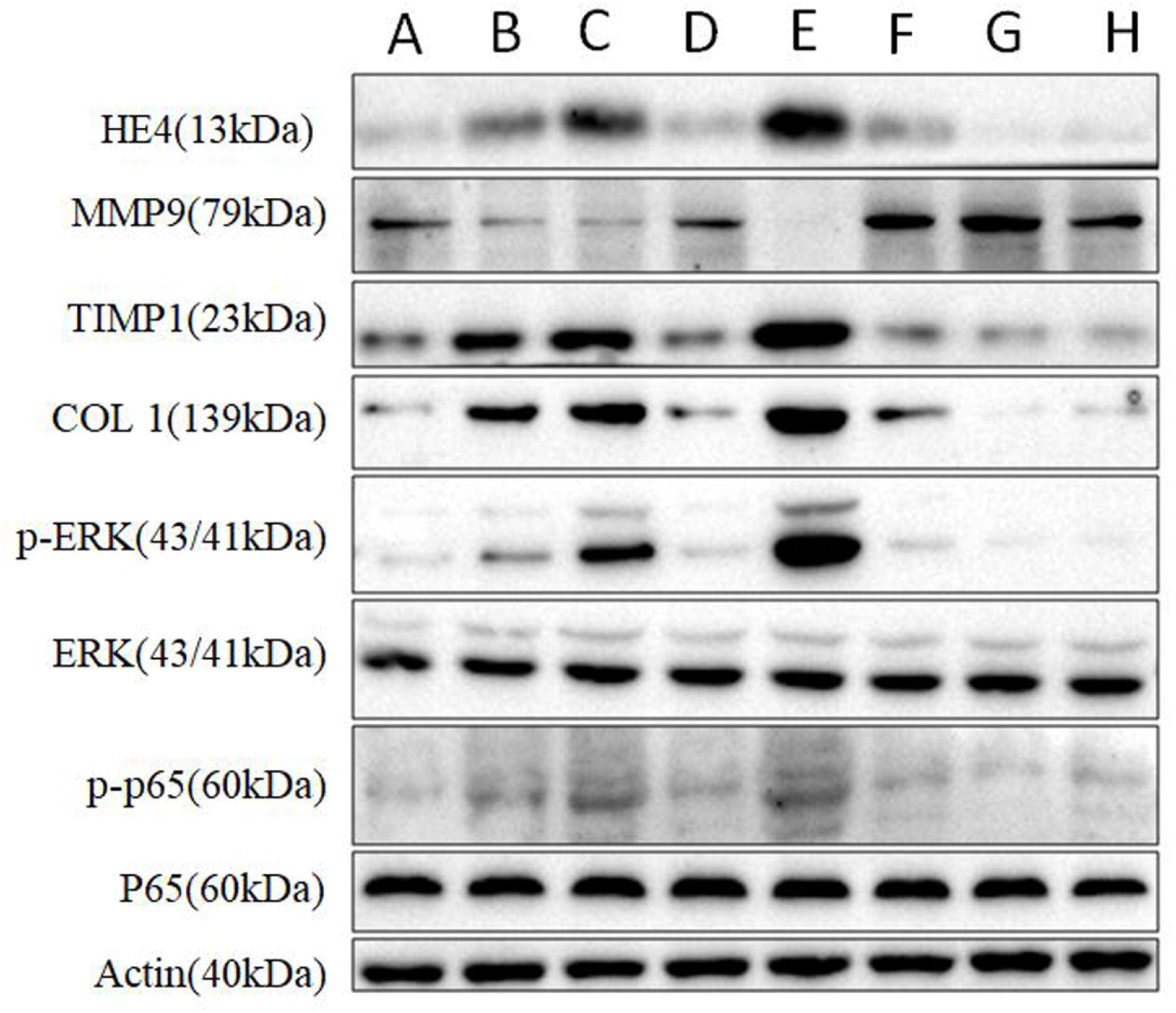
Target protein expression in cells of each group: A. normal control group; B: hyperoxia group; C: overexpression group; D: overexpression NC group; E: overexpression hyperoxia group; F: silent NC group; G: silent group; H: silent hyperoxia group (n=5)

### 6. RT-PCR detection of HE4, MMP9, TIMP1, COL1 mRNA expression in each group of cell samples

Compared with the normal control group, the expressions of HE4, TIMP1 and COL1 mRNA in the hyperoxia group were up-regulated, and the expression of MMP9 mRNA in the hyperoxia group was down-regulated compared with the normal control group, and the differences were statistically significant (p<0.05); Compared with the normal control group, the expression of MMP9 mRNA in the HE4 gene overexpression group was down-regulated, and the expression of TIMP1 and COL1 mRNA was up-regulated compared with the normal control group, and the differences were statistically significant (p<0.05); Compared with the normal control group, the expression of MMP9 mRNA in the gene silencing group was up-regulated (p<0.0.05); However, the mRNA expressions of TIMP1 and COL1 were similar to those in the normal control group, and the difference was not statistically significant (p>0.05).

The expression of MMP9 mRNA in the gene overexpression hyperoxia group was down-regulated in the high-oxygen group, and the expression of TIMP1 and COL1 mRNA in the high-oxygen group was up-regulated, and the differences were statistically significant (p<0.05); In the gene silencing hyperoxia group, the expression of MMP9 mRNA was significantly up-regulated in the hyperoxia group, while the expressions of TIMP1 and COL1 mRNA were down-regulated in the hyperoxia group, and the differences were statistically significant (p<0.05). (Figure 6).

**Figure 6:**
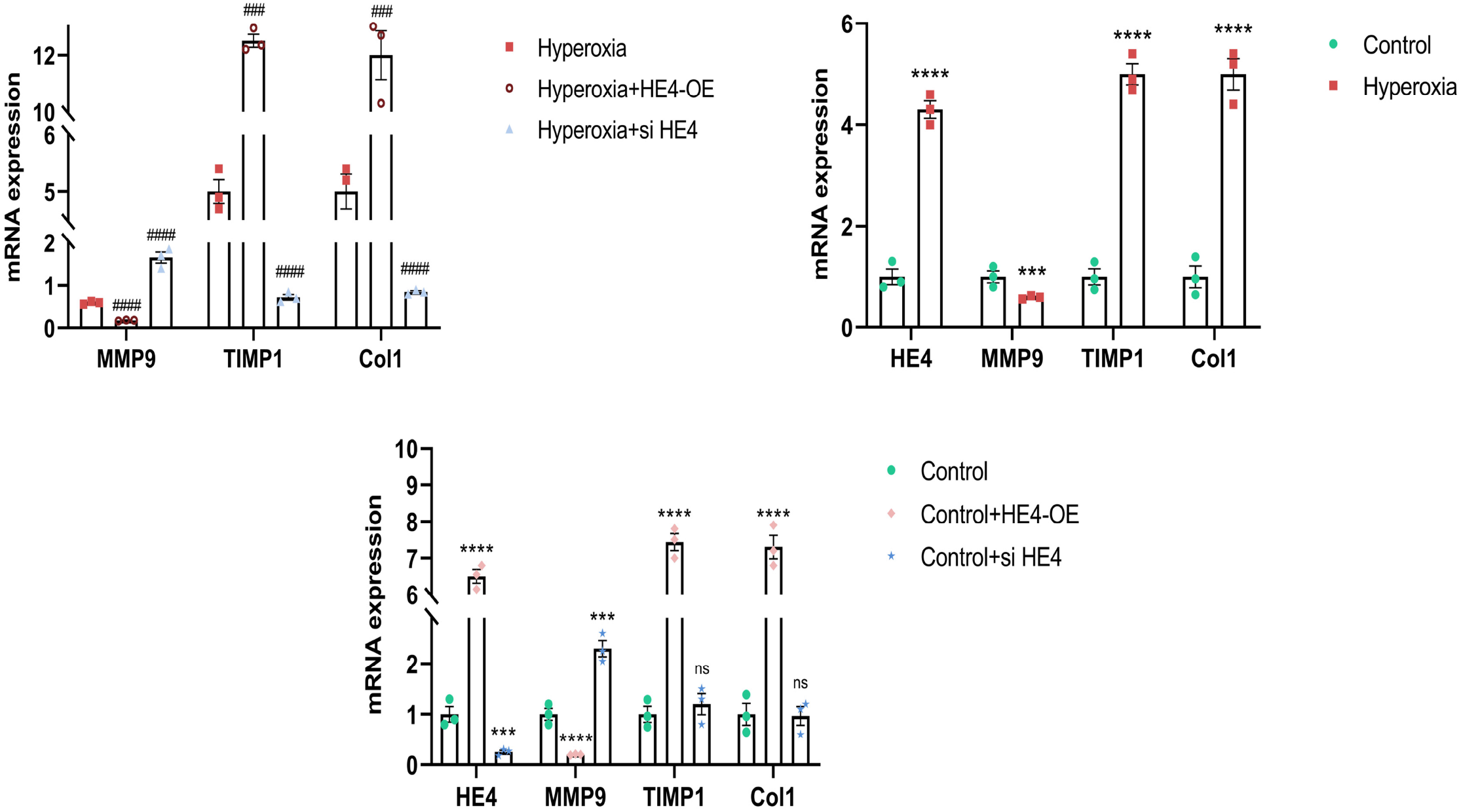
qPCR detection of mRNA expression in cell samples of each group (mean±SEM, n=3) ***p<0.001 and ****p<0.0001 vs. Control, ###p<0.001 and ####p0.0001 vs. Hyperoxia.

## Discussion

From the H&E staining light microscope, the alveolar septa of the hyperoxia-induced group were observed to be thin and discontinuous, the alveolar walls were thin, the alveolar cavity gradually increased, some alveoli were fused, the number of alveoli was reduced, the size was different, and the alveolar structure was simplified; The typical manifestations of alveolar simplification and alveolar cavity enlargement have appeared at 7 days of hyperoxia induction, which is consistent with the performance in various studies based on hyperoxia-induced BPD [11, 12]. In this experiment, hyperoxia intervention within 7 days resulted in acute lung injury in neonatal SD rats. The levels of HE4, MMP9 and TIMP1 in the serum of rats in the hyperoxia-induced group were significantly higher than those in the normal control group at 24 hours and 3 days. The protein content and mRNA transcription level were significantly higher than those of the normal control group at 3 days; However, they did not gradually increase with the prolongation of hyperoxia induction time and the aggravation of lung injury. HE4 is a matrix metalloproteinase inhibitor encoded by the WFDC2 gene. The high expression of HE4 mRNA can be detected in the trachea and lung [4], and it is highly restricted in normal airway epithelial cells. However, the expression increased when alveolar cells were injured, and the higher the level of HE4, the more severe the lung injury; For example, in the Nagy experiment, HE4 was highly expressed in the serum and lung biopsy tissue of patients with idiopathic fibrosis. A significant negative correlation [13, 14]; Zhang’s experiment also suggested that the level of HE4 is negatively correlated with lung function after interstitial lung injury in systemic sclerosis [15]; The higher the level of HE4, the worse the lung function; Raghu’s case-control study also found that serum HE4 was significantly elevated in patients with idiopathic pulmonary fibrosis (CF) [7]. In this experiment, the level of HE4 increased significantly when hyperoxia induced alveolar epithelial cell injury in rats. In the early stage of hyperoxia-induced acute lung injury, the level of HE4 increased with the progress of lung injury. The results of the early experiments were consistent, but HE4 It did not increase with worsening lung injury. In animal experiments, the protein expressions of MMP9 and TIMP1 in rat lung tissue and blood increased after hyperoxia intervention, and the change trend was inconsistent (MMP9 peaked at 24 hours after hyperoxia induction, and then gradually decreased, and TIMP1 was induced at 3 hours after hyperoxia induction. day is the peak). Matrix metalloproteinases (MMPs) and their tissue inhibitors (TIMPs) are involved in the whole process of acute lung injury and the degradation and remodeling of extracellular matrix in lung injury, and play an important role in the entire pathophysiological process of BPD[16–18]. In the physiological process of lung tissue, MMP9 and TIMP1 combine 1:1 to play a normal matrix remodeling effect. In acute lung injury, the expression of the two is abnormal, and the ratio of MMP9/TIMP1 is imbalanced. In the in vivo experiments, the expression of MMP9 and TIMP1 proteins was abnormal in hyperoxia-induced lung injury, and had different trends. The imbalance of protein ratios interfered with the normal remodeling process after lung injury.

From the in vivo experiments, it is known that there are changes in the gene expression and protein levels of HE4, MMP9 and TIMP1 in the lung tissue injury induced by hyperoxia. Both HE4 and MMP9/TIMP1 are significantly increased in the early stage of hyperoxia-induced acute lung injury. HE4 is also a recognized matrix metalloproteinase inhibitor, which can inhibit the activity of MMP9 and its activity of degrading collagen I (Collagen I, COLI) [3]; In hyperoxia-induced acute alveolar epithelial injury, MMP9 is a downstream molecule of NF-κB and ERK signaling pathways, which can directly degrade COLI and is inhibited by its natural inhibitor TIMP1. The imbalance between MMP9 and TIMP1 leads to abnormal basal remodeling. We speculate that HE4 should act on MMP9 and TIMP1 through a certain signaling pathway to mediate the process of lung injury remodeling when hyperoxia intervention causes alveolar damage; In order to further explore its mechanism of action, we selected primary rat alveolar type II epithelial cells as the research object, and carried out the overexpression and silencing of the HE4 gene during hyperoxia intervention, and selected a common type of hyperoxia-induced acute alveolar epithelial injury. NF-κB and ERK signaling molecules were studied to explore the possible mechanism of action.

In cell experiments, hyperoxia can induce the up-regulation of HE4 expression, accompanied by increased p-ERK and p-p65 protein contents, down-regulated downstream MMP9 expression, and increased COLI protein content and up-regulated expression; After HE4 gene overexpression, the protein contents of p-ERK and p-p65 were further increased, and the expression of MMP9 was further down-regulated. The ATII cells were further detected after HE4 gene silencing. Compared with the higher oxygen group, the protein contents of p-ERK and p-p65 were significantly decreased, the downstream MMP9 expression was up-regulated, and the COLI protein content was decreased and the expression was down-regulated. The changes of the above detection indexes after HE4 overexpression and hyperoxia intervention were more significant than those in the pure hyperoxia group. HE4 is a recognized matrix metalloproteinase inhibitor, which can inhibit the activity of MMP9 and inhibit its ability to degrade COLI [3]; it induces matrix remodeling and promotes collagen degradation through the ERK signaling pathway. In this experiment, the changes of the above detection indexes after HE4 overexpression and hyperoxia intervention were more significant than those in the pure hyperoxia group, suggesting that hyperoxia can cause high expression of HE4 in ATII cells, HE4 activates the phosphorylation of ERK and p65, and activates the downstream MMP9 Signaling pathway, inhibiting MMP9 mRNA expression, inhibiting protein activity, reducing COLI degradation, increasing collagen secretion, and promoting matrix remodeling and fibrosis, which are consistent with the results in in vitro experimental studies of renal fibrosis [3, 19]. In the experiment, the protein expression of p-p65/P65, a key protein of NF-κB signaling pathway induced by hyperoxia, was lower than that of p-ERK/ERK. Both HE4 overexpression and hyperoxia induction can promote the phosphorylation of p65 and aggravate hyperoxia-induced alveolar cell injury; Compared with the control group, p-p65 in the hyperoxia group and HE4 overexpression group was higher than that in the control group. Similar findings were found in Wang’s experimental study [20], but the protein expression of p-p65/P65 was higher than that of p-ERK/ERK. It should be considered that the experiment only uses hyperoxia as an interfering factor, and the action time is 6 hours. The early damage caused by hyperoxia is mainly related to the damage caused by reactive oxygen species, and is mainly related to the ERK signaling pathway [21, 22].

MMP9 is co-secreted by alveolar epithelium and alveolar macrophages. TIMP1 is a natural inhibitor of MMP9, inhibiting the activity of MMP9, inhibiting the degradation of collagen, and maintaining the integrity of the extracellular matrix. The TIMP1 protein has two distinct domains: an N-terminal domain and a C-terminal domain. The N-terminal domain can form a non-covalent complex with MMP9 at a ratio of 1:1 to inhibit the activity of MMP9, inhibit the degradation of collagen, and maintain the integrity of the extracellular matrix [23]; The C-terminal domain can play a signal transduction role by binding to CD63, CD82, etc. [24, 25]; TIMP1/MMP9 has three possible proteolytic states, which affect the signal transduction ability of TIMP1. In Thevenard’s experiment, the amount of MMP9 protein was decreased, and the protein content of TIMP1 was increased. TIMP1 combined with CD82, CD63 and LPR1 through the C-terminal domain to play a signal transduction role [26], inducing intracellular signaling pathways, which further inhibited the activity of MMP9. produce regulation. In this experiment, the expression of TIMP1 was up-regulated in the hyperoxia group, and the expression of TIMP1 mRNA was significantly up-regulated after HE4 gene overexpression compared with the normal control group, and the up-regulation was more significant after hyperoxia and gene overexpression. resemblance; The expression of TIMP1 mRNA in the gene silencing group was similar to that in the normal control group, and the expression of TIMP1 was significantly down-regulated after gene silencing under high oxygen conditions. HE4 can activate the downstream MMP9/TIMP1 signaling pathway through the ERK and NF-B signaling pathways, inhibit the expression of MMP9 mRNA, and reduce protein synthesis; The expression of TIMP1 mRNA is up-regulated, and the protein synthesis is increased; the increased content of TIMP1 cannot bind with MMP9 in a 1:1 non-covalent manner, and the ratio is unbalanced to promote collagen degradation, COLI is increased, and the basal remodeling function is disordered; In addition, the free C-terminal domain of TIMP1 can mediate basal remodeling through other signaling pathways [27]. In this experiment, the mRNA expressions of HE4 and TIMP1 were both up-regulated, the protein synthesis was increased, and the protein contents of p-ERK and p-p65 were increased. p-p65, and then the expression of TIMP1 mRNA up-regulated and inhibited the activity of MMP9. Similar results were obtained in Zhang’s experiment, that is, HE4 overexpression in HK2 cell line inhibits ECM degradation through P65 phosphorylation and nuclear translocation to activate the NF-κB pathway, upregulate TIMP1, and inhibit the activity of matrix metalloproteinases [28]. It should also be considered that HE4 can directly regulate TIMP1 through a signaling pathway in alveolar epithelial cell injury.

Hyperoxia can induce ATII cells to overexpress HE4, promote p-ERK and p-p65 phosphorylation to activate downstream signaling pathways, inhibit MMP9 expression, reduce COLI degradation, increase collagen secretion, and promote matrix remodeling; HE4 cooperated with TIMP1 to inhibit the activity of MMP9 to a certain extent. However, both HE4 and TIMP1 are inhibitors of MMP9. Why are they elevated at the same time in in vitro cell experiments, and the mechanism of action of HE4 and TIMP1 is still unclear, and further research is needed.

### Conclusion

In conclusion, this study used hyperoxia-induced neonatal SD rats to discover the mechanism of HE4 in alveolar damage. Through overexpression and gene silencing of HE4 in vitro experiments, we found that HE4 mediates the pathophysiological process of hyperoxia-induced lung injury in rats through ERK, MMP9, and TIMP1 signaling pathways. HE4 overexpression mediates lung basal remodeling and lung injury.

## Notes

### Competing Interest Statement

The authors have declared no competing interest.

